# Colormesh: A novel method for quantifying variation in complex color patterns

**DOI:** 10.1101/2020.07.17.205369

**Authors:** Jennifer J. Valvo, F. Helen Rodd, David Houle, J. David Aponte, Mitchel J. Daniel, Kenna Dwinell, Kimberly A. Hughes

## Abstract

Color variation is one of the most obvious examples of variation in nature. Objective quantification and interpretation of variation in color and complex patterns is challenging. Assessment of variation in color patterns is limited by the reduction of color into categorical measures and lack of spatial information. We present Colormesh as a novel method for analyzing complex color patterns that offers unique capabilities. Compared to other methods, Colormesh maintains the continuous measure of color at individual sampling points throughout the pattern. This is particularly useful for analyses of variation in color patterns, whether interest is in specific locations or the pattern as a whole. In our approach, the use of Delaunay triangulation to determine sampling location eliminates the need for color patterns to have clearly defined pattern elements, and users are not required to identify discrete color categories. This method is complementary to several other methods available for color pattern quantification, and can be usefully deployed to address a wide range of questions about color pattern variation.

## Introduction

Variation in color is one of the most prolific and obvious examples of variation in nature. Measurement of color is used in many fields of study including, for example, astronomy (Bessell 2005, Robinson et al. 2010), medicine (Bhargava and Madabhushi 2016, Limkin et al. 2017), and agriculture and food science (Dell’Aquila 2009, Pathare et al. 2013). In biology, ecological, evolutionary, and mechanistic investigations of organismal color have been prominent in the scientific literature since the 19^th^ century and have provided key insights into many aspects of organismal function and evolution, such as the role of coloration in thermoregulation, crypsis, aposematism, mimicry, mate choice, and speciation (Darwin 1859, Cott 1940, Jennions and Petrie 1997, Mallet and Joron 1999, Mappes et al. 2005, Mclean and Stuart-Fox 2014, Cuthill 2019). In recent years, color analyses have expanded to include a more intricate understanding of how both color and patterns are produced (e.g., Manukyan et al. 2017, Shawkey and D’Alba 2017) and how they are perceived (e.g., Stoddard et al. 2014, Kelber 2016) allowing for novel questions to be asked about the role of organismal coloration.

Despite the importance of color, in many taxa the complexity of color and its patterning pose serious challenges to measurement and interpretation. Many different approaches, that often rely on human perception of color, have been used to quantify color patterns and how they vary among organisms. Early methods of assessing color in biology relied on categorical schemes, such as assigning patterns to discrete morphs (Tan and Li 1934, Semler 1971, Brown and Clegg 1984). Later studies quantified color using measures of total area or percent coverage of discrete categories of color (e.g., “orange”, “blue”, etc.), as well as the frequency of particular color pattern elements (e.g., numbers of “spots” of particular colors) (Houde 1987, Petrie and Halliday 1994, Olsson 1994). These relatively low-dimensional measures were and are useful in addressing biological questions such as: Is there covariation between female preference and ornamental traits (Endler and Houde 1995, Ellers and Boggs 2003)? Are some color morphs more successful at attracting mates or surviving (Petrie 1992, Sinervo and Lively 1996, Borer et al. 2010)? Do morphs differ in life-history traits (Svensson et al. 2001, Emaresi et al. 2014)? Does morph frequency vary with habitat type (Power et al. 2005, Ahnesjö and Forsman 2006)?

In these studies, human perception was used to determine color categories and the boundaries of color pattern elements in order to quantify color patterns. Recent technological advancements in digital imaging and computation have enabled more objective characterization of these metrics. For example, individual pixels in digital images can be assigned values in a color space (e.g., red, green, and blue color channels in RGB color space), and spectrophotometry can capture the entire reflectance spectra of specific locations on an organism. In digital image analysis, every pixel can be described by a quantitative value for each color channel. Pixels can then be grouped into discrete color categories using clustering or thresholding to perform image segmentation. Image segmentation is easily performed by software such as ImageJ (Schneider et al. 2012) for use with cell counting and counting pixels within a specified color range to identify diseased tissue (Papadopulos et al. 2007, Hadi et al. 2011, Schindelin et al. 2015). Packages within the R statistical computing environment (R Core Team 2019) offer several methods for grouping similar colors to assess overall patterns. For example, the *patternize* package (Van Belleghem et al. 2018) can be used to compare the similarity of discrete color categories, while *colordistance* (Weller and Westneat 2019) can evaluate color category differences in color space.

In color segmentation approaches, color categories require user determined thresholds which introduces a potentially problematic level of human subjectivity (Davidoff and Fagot 2010, Siuda-Krzywicka et al. 2019; but see Bergeron and Fuller 2018). Reflectance spectrometry is an objective measure of color that measures the wavelengths and intensities of light reflected from a small point sample over a continuous range of wavelengths (Endler 1990, Zuk and Decruyenaere 1994, Andersson et al. 1998, Gomez and Théry 2007). Heterogeneous patterns can be sampled by collecting data from multiple sampling points in a standardized manner (Cuthill et al. 1999, Endler and Mielke 2005, Endler 2012). This technique has been used to acquire objective color data in many ecological and evolutionary studies, often in combination with models of the visual sensitivity of receivers of the color information, e.g., potential predators, mates, or pollinators (Cortesi and Cheney 2010, Stoddard and Stevens 2011, Dyer et al. 2012, Isaac and Gregory 2013). Understanding how a receiver perceives a visual signal is clearly important for studies of behavioral and ecological interactions based on color, and several approaches have been developed to make use of this information (e.g. Endler and Mielke 2005, Endler 2012, Endler et al. 2018, Maia et al. 2019). However, because a large number of spectrometric sample points are required when color patterns are complex, using this approach is not currently feasible for comparing complexity of entire color patterns among a large number of individuals.

Here we present Colormesh, a new method for quantifying color patterns from digital images that is objective, high dimensional, high throughput, and that does not rely on threshold setting or subjective determination of the number or type of color categories. Colormesh captures multi-dimensional color data by dense sampling of quantitative color values across the entire sample area. To accomplish this, our method standardizes images using geometric morphometrics and identifies homologous sampling points using Delaunay triangulation. This novel use of Delaunay triangulation allows us to address biological questions without *a priori* assumptions about color categories or the spatial properties of the color pattern. With Colormesh, we enable (1) the analysis of color patterns that are spatially complex and/or lack well-organized color pattern elements (e.g., spots or stripes), (2) the capture the continuous high-dimensional nature of color, (3) the objective sampling of color values from standard digital images, with straightforward extension to wavelengths outside the human-visible range, and (4) flexibility in color sampling density and the size of the area sampled.

We demonstrate the utility of the Colormesh pipeline using color data from eight natural and three experimental populations (described below) of the Trinidadian guppy (*Poecilia reticulata*), a model system in which the evolution of color has frequently been studied. Complex color patterns in this species are male limited, highly heritable, and highly variable both within and between populations (Winge 1927, Endler and Houde 1995, Houde 1997, Hughes et al. 1999, 2013, Brooks and Endler 2001b, Magurran 2005, Olendorf et al. 2006, Kemp et al. 2008, Gordon et al. 2015a). Male guppy coloration has been the focus for studies of local adaptation (Houde 1997, P. Millar et al. 2006, Gordon et al. 2015b) and for the maintenance of ecologically-important variation (Hughes et al. 1999, 2013, Brooks and Endler 2001a, Olendorf et al. 2006, Evans et al. 2008, Valvo et al. 2019). Guppies have repeatedly evolved in response to ecological conditions above and below barrier waterfalls in Trinidadian rivers and streams (Endler 1978, Magurran 2005), and upstream populations are known to be descendants of downstream populations within rivers (Magurran et al. 1992, Shaw et al. 1992, Becher and Magurran 2000, Alexander et al. 2006, Crispo et al. 2006, Suk and Neff 2009, Willing et al. 2010). This repeated ecological transition has led to extensive parallel evolution of many phenotypes (Reznick and Endler 1982, Reznick and Bryga 1987, Reznick 1989, Reznick et al. 1996b, Reznick and Bryga 1996, Rodd and Reznick 1997, Torres Dowdall et al. 2012), including male color (Endler 1978, 1983, 1991). Ecological differences between down- and upstream sites such as predation have also been proposed to select for differing levels of color polymorphism between populations (Endler 1980, Olendorf et al. 2006, Fraser et al. 2013). Here we have used Colormesh to address three questions about color variation and evolution in Trinidadian guppies. We first asked if our method could successfully classify samples into the correct population. We then asked if the (multivariate) direction of color evolution between up- and downstream populations was consistent, or parallel, in different river drainages. Finally, we asked if the within-population variance in color differed consistently between up- and downstream populations, and evaluated the spatial distribution of within-population variance.

## Methods

### Overview

Each section below provides information detailing the Colormesh pipeline. We first describe a general pipeline for processing images using geometric morphometrics to standardize images, which allows for the sampling of homologous locations among images. We then describe how color is sampled and some of the built-in flexibility of the pipeline for the sampling process. Finally, we apply Colormesh to the Trinidadian guppy system to demonstrate applications of the color pattern analysis pipeline to evolutionary questions relevant to the study system.

### The Colormesh pipeline

#### Image processing

Digital photography is used to capture images of specimens that also include a size scale and color standard. Digital images are then processed using the *TPS Series* morphometrics software (Rohlf 2015); this software must be used within a Windows operating system. The *tspUtil* (V. 1.78) software is a utility program used to prepare files for further processing in the *TPS series* (Rohlf 2015). Using *tspUtil*, a text file (.TPS) containing the file path for each image is first created to prepare the images for landmark placement. The *tpsDig2* (V. 2.31) software (Rohlf 2017) is then used to open the image, set the scale, and place landmarks for shape analysis. As landmarks are placed, the TPS file is populated with the x, y coordinates of landmarks specific to each photograph. Initially, traditional landmarks are placed at several locations on the specimen. Placement of traditional landmarks at locations that can be consistently identified across samples (e.g., fin attachment sites) facilitates comparisons of samples that differ in size and shape. The user then has the option to place semilandmarks (sliding landmarks), approximately equally spaced, between the traditional landmarks. Semilandmarks are placed to represent a curve or surface where locations that are biologically homologous are not easily identifiable (Bookstein 1997); the number of semilandmarks depends on the complexity and level of variation of the curve (Gunz and Mitteroecker 2013). Sliding of semilandmarks reduces shape variation due to unequal distribution of semilandmark placement between the traditional landmarks. All landmarks should be placed in a consistent order among each photo.

The next step in image processing is to generate a consensus (average) shape of specimens using the TPS file now populated with x,y coordinates of landmarks. Using the “make sliders file” operation within the *tspUtil* software, semilandmarks are first identified and stored as a separate file (.NTS). The *tpsRelw32* (v. 1.70) software is then used to compute the consensus specimen shape (Rohlf 2018) using the option to let semilandmarks slide to minimize bending energy between individuals and the consensus shape (Gunz and Mitteroecker 2013). The consensus shape is calculated by performing a least-squares orthogonal generalized Procrustes analysis using both the NTS and the TPS files as inputs. The resulting landmark coordinates of this overall consensus shape are saved as a new TPS file to be used as the target for the last step in image processing.

The final step in image processing is to “unwarp” images using *tpsSuper* (V. 2.05) (Rohlf 2015). This process is necessary because it transformed each specimen to have the same shape to account for size and shape differences among individuals. The unwarping process transforms each subject to the consensus shape by mapping pixels from the consensus shape back to the corresponding pixels in the original image. Mapping pixels in this direction ensures all pixels from the original image are represented in the image that has been transformed to the consensus shape (Rohlf 2015). Because each image is unwarped to the same consensus shape and pixels are mapped from the consensus shape back to the original image, the unwarped images have the same pixel dimensions. At the completion of the unwarping process in *tpsSuper*, the unwarped image files (TIFs) are saved to be sampled for color as described below. It is important to note that the unwarping process is computer memory intensive; the maximum number of photos the *tpsSuper* software can process at one time depends on available computer RAM, the number of images being unwarped, and image size. If *tpsSuper* is unable to process the all the images included in the original TPS file, the user can edit the TPS file in a text editor (e.g., Notepad) and break them into smaller groups for unwarping to the target consensus shape.

A separate TPS file is needed for the color calibration process (described below) since these landmark coordinates are solely used to identify sampling locations on the color standard. A duplicate TPS file is created with the *tspUtil* software to identify coordinates to sample on the color standard. Using the *tpsDig2* software to open this duplicate TPS file, a landmark is placed on each color of the standard included in each image, providing image-specific locations to sample for the calibration process; the order of landmark placement on colors of the standard should be consistent among images.

#### Color sampling and calibration

The Colormesh pipeline uses Delaunay triangulation to identify comparable pixels on each specimen that will be sampled for RGB values (see Aurenhammer 1991 for use in geometric data structure). Delaunay triangulation is used in surface reconstruction where a large number of points within a boundary surface (i.e., a finite set of points) are reduced to a concise representation of the shape (De Berg et al. 2008, Bala and Sekhon 2011). Custom functions in the R statistical computing environment V. 3.5.3 (R Core Team 2019) were developed to collect color data at sample locations determined using Delaunay triangulation. All color sampling and calibration processes were performed in R (R Core Team 2019).

The first step in the pipeline is to read in the landmark coordinates for the overall consensus shape using the *read.tps* function (https://gist.github.com/mrdwab/2062329). A surface of points is then distributed within the overall consensus shape using a custom function (*tri.surf*). The first round of triangulation uses the x, y coordinates of the landmarks as the vertices (nodes). The *tri.surf* function then calculates the coordinates of the centroid for each triangle, and stores the resulting x, y coordinates in a matrix. If additional rounds of triangulation are specified by the user, the *tri.surf* function uses the saved x, y coordinates of the centroids from the previous round of triangulation as the vertices for the subsequent round. Each additional round of triangulation therefore increases the density of sampling points allowing for user controlled granularity of color sampling (e.g., Figure 2A). However, the increase in sample size also increases the dimensionality of the data and therefore downstream processing times will likely increase as well. An approach for deciding the number of rounds of Delaunay triangulation (i.e., density of sampling) used in later applications is described below (see the Multivariate classification and differentiation among populations section).

In addition to the density of sampling, the user also defines the size of the sampling circle using the custom *sampling.circle* function. This function calculates the average values for each color channel (i.e., R, G, and B) for all pixels within the sampling circle (e.g., Figure 2B). The maximum size of the sampling circle is limited to avoid overlapping circles and thus resampling pixels; therefore, the sampling circle size limit depends on the density of sampling points. The user-determined size of the sampling circle allows the user to control the level of pixel averaging for color sampled at a location. Sample circle sizes are defined by the radius measured in pixels. In contrast to the density of sampling points, the sampling circle size does not influence downstream processing time.

To sample color, the *readImage* function in the R package *EBImage* (Pau et al. 2010) is used to extract R, G, and B values at the x,y coordinates identified by the *tri.surf* function (above) from each unwarped image (e.g., Figure 2C & D). The extracted R, G, and B values for each sample location are then stored in matrices specific to each color channel; when sampling circles are greater than 1 pixel, the mean values for all pixels within each sampling circle are stored.

To calibrate the RGB values sampled from each image, a color channel correction is calculated based on the known RGB values from the color standard. The *read.tps* function is first used to read in the duplicate TPS file containing the coordinates of the landmarks placed on the color standard. The *readImage* function samples the RGB values of the pixels contained within a five-pixel diameter sampling circle centered on the coordinates provided by the TPS file; the user has the option to change the sampling circle size. Similar to the image sampling described above, a separate matrix is created for each color channel. Thus, three matrices are generated, each having rows equal to the number of photos and columns equal to the number of landmarks placed on the color standard. To determine the image-specific color correction, a color channel correction vector is then generated by calculating the mean deviation from the known color channel values of the color standard included in each image. The color channel specific correction vector therefore has a length equal to the number of images; each image has a unique correction for that color channel. The correction vector for each color channel is then applied to matrices containing values sampled from the unwarped specimen images. Because RGB color data extracted using *EBimage* is on a scale from 0 to 1, corrected values are limited to this range by default.

Each record then consists of a color vector with *k* points * three (or more) color channel values. For example, in the application of the Colormesh pipeline described below, an image sampled using four Delaunay triangulations had a vector with 7386 values (2462 sample points * 3 color channels). If data from additional color channels (e.g., ultraviolet) are available, these data are easily added to the data structure to provide an additional 2462 dimensions.

### Colormesh applied to the Trinidadian guppy system

#### Populations sampled

To compare color patterns in a diverse set of populations, we photographed wild male guppies in May of 2016 and 2017. These fish were collected from six rivers and, in some cases, their tributaries belonging to three major drainage systems in the Northern Range in Trinidad (see Supplemental Table S1 for GPS locations): the Caroni drainage (Aripo, El Cedro, and Guanapo Rivers), Northern drainage (Marianne and Paria Rivers), and the Oropuche drainage (Turure River). In the Aripo, El Cedro, Guanapo, Marianne and Turure Rivers, we sampled males from up- and downstream habitats that are characterized by important ecological differences (Houde 1997, Rodd et al. 2002, Magurran 2005). Downstream habitats are often referred to as high-predation due to the presence of one or more species of large piscivorous fish (Endler 1980, Reznick et al. 1996a). In addition, many downstream sites have relatively open forest canopy and high primary productivity (Grether et al. 2001). In contrast, upstream habitats, typically referred to as low-predation, are located above a barrier waterfall and generally contain one main, smaller guppy predator (*Rivulus hartii*) that preys on juveniles and small adult guppies (Gilliam et al. 1993, Reznick et al. 1996b). These low-predation habitats typically have relatively closed canopy and low primary productivity (Grether et al. 2001, Reznick et al. 2001). We also sampled one additional population, the Houde Tributary of the Paria River, which is considered low-predation and does not have an associated high-predation site (Houde 1997, Magurran 2005). Of the 11 populations sampled, eight were naturally-occurring populations and three were introduced populations. The high-and low-predation populations sampled from the Turure river were originally high-predation fish from the Arima river (Caroni drainage system) that were introduced to the upstream Turure river (Oropuche drainage system) in 1957 (Becher and Magurran 2000). Fish sampled from the low-predation El Cedro population were also descendants from an introduction experiment where high-predation fish were transplanted upstream within the river (Reznick and Bryga 1987).

For logistical reasons, we collected and photographed the fish in two different years, both in the dry season. Males from the Aripo, El Cedro, and Paria Rivers, and the low-predation population of the Marianne River were collected and photographed in 2016. Males from the Guanapo, Turure, and the high-predation population of the Marianne River were photographed in 2017. For each of the 11 populations, we sampled 24-57 males per population (See Supplemental Table 1 for number of males sampled per population).

#### Ethical note

All field methods and protocols were approved by the Florida State University Animal Care and Use Committee (protocol No. 1442 approved on 29 October 2014 and protocol No. 1740 approved on 16 October 2017). Research permits were granted by the Ministry of Agriculture, Land, and Fisheries (Aquaculture Unit) of the Republic of Trinidad and Tobago in March of 2016 and May of 2017. Fish were caught using butterfly nets and transported in sealed Nalgene bottles containing water with Stress Coat (API) to the nearby William Beebe Tropical Research Station, located in the lower Arima valley in the Northern Range, Trinidad. Fish were separated by population and maintained in single-sex 20-40L aquaria for 24-48 hours prior to taking their photographs. We collected digital photographs of male guppies by lightly anesthetizing them with buffered MS222. A camel-hair paintbrush was used to position the dorsal fin and gonopodium away from the body of each fish. Fish bodies were dabbed with a Kimwipe to remove excess water and reduce glare. Water exchanges of 30% were performed daily using conditioned rainwater collected on site. Fish were fed Tetramin Tropical flake food twice daily.

#### Image collection and processing

To compare color patterns, we collected digital photographs of male guppies (N=485) from the 11 populations described above (see Supplemental Methods for details on fish and image collection). A size and color standard were included in each image. All data processing steps were performed on a Dell Precision with an Intel i7 CPU, Windows 10, and 16GB RAM. Images were processed using the *TPS Series* morphometrics software as described above. Because the final step in processing images uses computer memory to determine the placement of pixels in the unwarped images, we first reduced the number of pixels in each photo by cropping pixels from the area surrounding the fish. To do this, we used the batch cropping ability in Photoshop CC 2018. Photo size was reduced from 4368 × 2912 pixels to approximately 2000 × 1000 pixels for images taken in 2016 and 1350 × 400 pixels for 2017 photos; differences in photo dimension within each batch depended on minor variation in placement of fish within photos. Because cropping removed the scale and color standard from the image, we created a duplicate set of images for setting the scale and calibrating color (see below).

We first used the *tpsUtil* (V.1.78) software to build a separate TPS file for each of our populations’ image sets (cropped and original). Using the *tpsDig2* (V. 2.31) software, a lab assistant (K.D.) placed 62 landmarks around the perimeter of each fish within the set of cropped images. Seven traditional landmarks were first placed at the following locations: the tip of the snout, anterior and posterior connection points of the dorsal fin to the body, dorsal and ventral connection points of the caudal fin to the caudal peduncle, posterior and anterior connection points of the gonopodium to the body (Figure 1). The observer then placed 55 additional semilandmarks among the traditional landmarks; all landmarks were placed in a counterclockwise direction (Figure 1). Using the original uncropped image sets, landmarks were placed on each of the five colors in the color standard for use in calibration. To include the size standard with the appropriate TPS files for the cropped images, we used *tpsDig2* to calculate the image scale (pixels/cm) from the original uncropped images and added the scale information to each TPS file containing cropped images.

**Figure 1.**
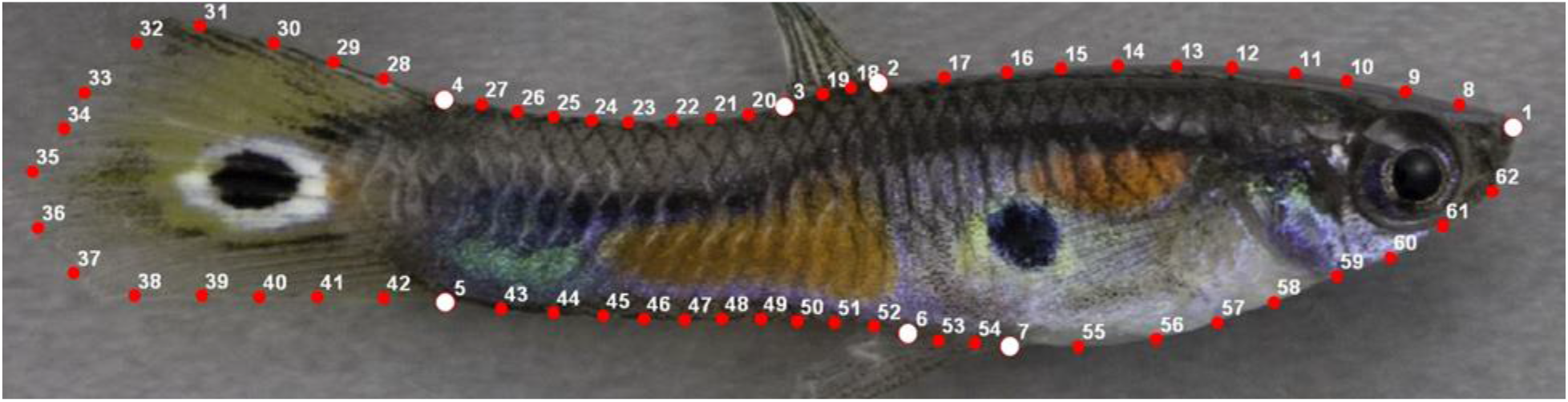
An example of an image opened within the *tpsDig2* software that was used to place landmarks on an image. A total of 62 landmarks are placed in the same order on every image. Seven traditional landmarks (white circles) are placed first in locations that can be consistently identified on each fish. Between the traditional landmarks, 55 semilandmarks (red circles) are placed; the quantity between each traditional landmark is consistent between images and the user placed them approximately equally spaced apart.

Populations were initially processed separately to generate population-specific consensus shapes using the *tpsRelw32* (v. 1.70) software (Rohlf 2018). Population-specific consensus shapes were determined prior to the overall consensus shape to avoid biasing fish shape toward a particular population since the number of fish sampled for populations varied. We saved the resulting landmark coordinates of the population-specific consensus shape into a new TPS file. To generate a consensus shape for all populations, we repeated this process using the TPS file we created that contained the landmark coordinates of the population-specific consensus shapes. The landmark coordinates of this overall consensus shape were then used as the target in the final step of image processing where images are unwarped to this shape. All cropped images were then unwarped to overall consensus shape using *tpsSuper* (V. 2.05) and saved as TIF files.

Using the Colormesh pipeline (described above), RGB color values were then extracted from each unwarped fish image. To calibrate these values, color corrections were applied based on color values extracted from the color standard in the original images. Because the known RGB values of the five colors included on the color standard were given on a scale of 0 to 255, values were divided by 255 to match the scale of values extracted by *EBImage* which range from 0 to 1 (Supplemental Table S2).

To evaluate the consequences of using different sampling schemes we specified two, three, and four triangulations (Figure 2A) to provide increasingly higher sampling density of the image (302, 842, and 2486 points, respectively). For each of these sampling densities, we evaluated four sizes of sampling circle (1, 3, 5, and 9 pixels in diameter, Figure 2B) to provide an increasing degree of pixel averaging. The higher the level of sampling, the more precise the representation of the image (Figure 2C & D). Changes in sampling circle diameter did not produce a noticeable difference in image representation (see Supplemental Figure S3).

**Figure 2.**
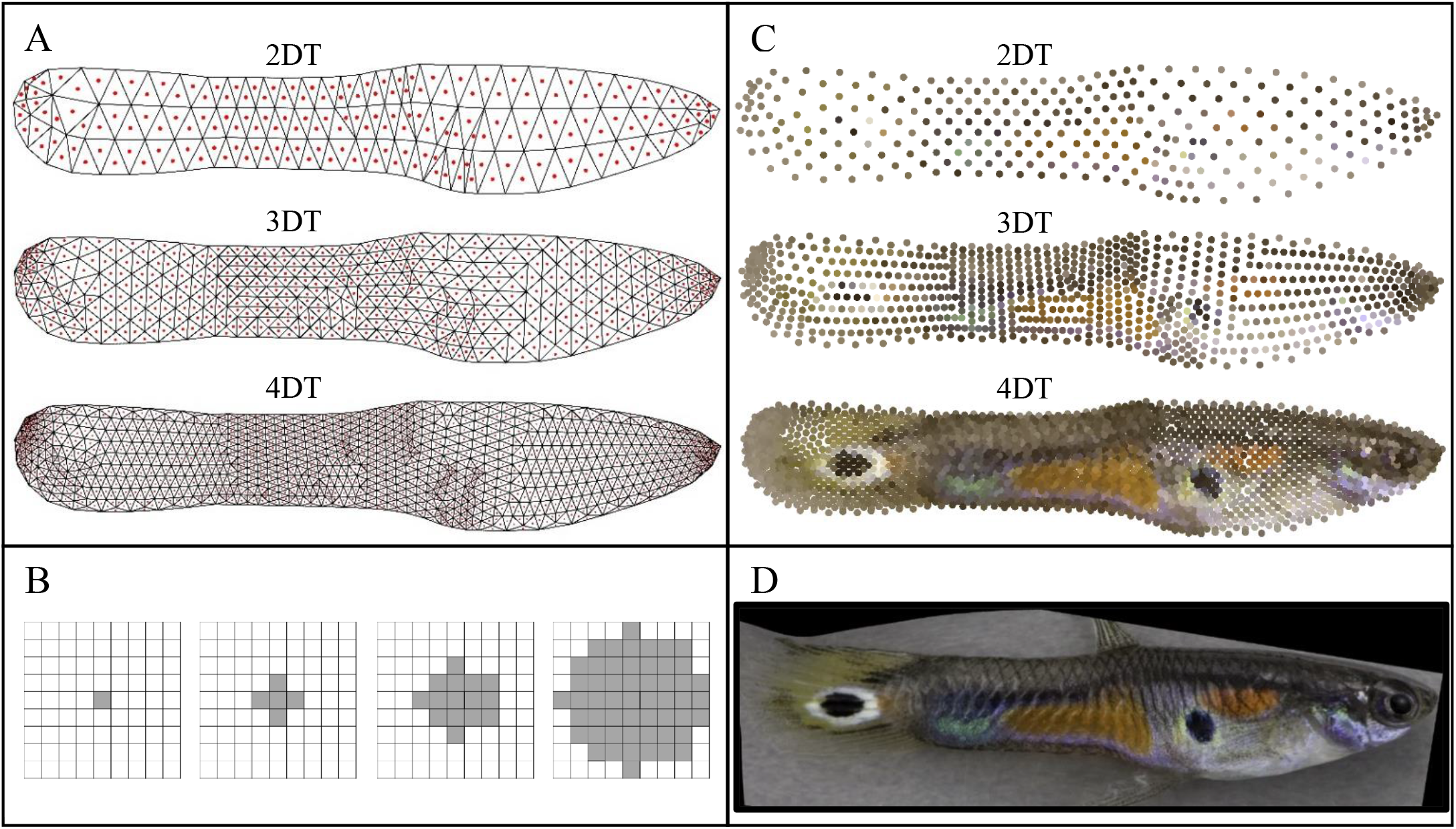
Panel (A) shows the mesh generated from the user-defined sampling scheme of 2, 3, and 4 rounds of Delaunay triangulation (DT). The triangle mesh is used to determine sampling location (red dot at each triangle centroid). The triangle centroids defined by the previous round of triangulation were used to generate the subsequent round of triangulation. Panel (B) shows the pixels sampled depending on the user-defined sampling circle radius size. Shown are the sampling circles with diameter = 1, 3, 5, and 9 pixels, from left to right. Panel (C) shows a plot of the RGB values sampled from an unwarped image (Panel D) at different sampling densities (DT’s). Here we only show the sampling design where the sampling circle diameter =1 (only the triangle centroid RGB values were measured). To visualize the color sampled at single pixels, the sampled RGB values from each pixel are reproduced as larger circles.

### Multivariate classification and differentiation among populations

When individuals are known to belong to particular groups *a priori*, Discriminant Analysis of Principal Components (DAPC) is often used on high dimensional data sets to describe the properties of those groups. DAPC was first proposed for analysis of genetic marker data by Jombart et al. (2010). In the R package, *adegenet* (Jombart 2008), the *dapc* function performs a Principal Components Analysis (PCA) followed by a discriminant analysis to define the linear combinations of principal component scores that minimize within and maximize between group variances.

DAPC can also be used in a cross-validation framework to determine if data can be used to classify observations into pre-defined categories. We used DAPC to determine which of the sampling schemes and sample circle sizes (described above) captured RGB color data that allowed us to best discriminate among populations. Because DAPC requires the user to specify how many PC’s to retain, we first used the *xvalDapc* function in the R package *adegenet* (Jombart 2008) on each of the calibrated RGB datasets (three sampling schemes * four sample circle sizes = 12 datasets total). This cross-validation function performs a DAPC on a training and validation set of data and reports the proportion of successful assignments and root mean squared error (RMSE) for a varying number of retained PC’s. For our cross-validation, we defined the training and validation populations as 80% and 20%, respectively; therefore, data from 80% of the samples in each of the 11 populations were used as training data to fit functions for the Discriminant Analysis and the remaining 20% of data were used to evaluate the prediction success of the functions. We defined the maximum number of PC’s to retain during the DAPC cross-validation as (*n.pca*) = 485 (the number of fish images). Thus cross-validations were performed at nine different retention levels of PC’s in increments of 50 ranging from 50 to 450. We set the number of replicates to be carried out at each PC retention (*n.rep*) = 100. We compared the lowest RMSE and proportion of successful placements for each of the 12 sampling schemes to determine which sampling scheme would be used for the remaining analyses.

### Direction of evolution within and between rivers

We used the color data collected by Colormesh to compare the direction of evolutionary divergence between populations with known ancestor-descendant relationships. This analysis tests the hypothesis that color pattern evolution has evolved convergently each time high-predation guppies have invaded low-predation habitats. For four of the six rivers that we sampled (Aripo, El Cedro, Guanapo, and Turure), the paired high- and low-predation populations were sampled in the same year. This meant the images had the same pixel resolution prior to the unwarping process described above, so the high- and low-predation populations within these four rivers could be compared directly. For this analysis, we used the RGB color data set collected from the four Delaunay triangulation sampling scheme and a sample circle diameter = 1. With this sampling scheme, each fish had R, G, and B values measured at 2462 positions for a color vector of length = 7386 variables.

To produce an average color pattern for a given population, we calculated the mean value of each color channel at each sampling point. Thus, each population had a mean color vector consisting of 2462 values for each color channel. Because low-predation populations are known to be descendants of high-predation populations within each river, the difference between the population mean color vectors measures the direction of color evolution within a river. If color pattern evolution was similar among rivers, these difference-vectors should be more similar than that expected by chance. We measured the similarity of these vectors indicating the directions of color evolution by calculating the angle between the vectors of each pair of rivers. If colors evolved similarly in two river drainages, the vectors indicating direction of evolution will have a smaller angle between them than random vectors. The angle between the vectors can also be expressed as a vector correlation (*r*). This is mathematically equivalent to Pearson’s correlation coefficient.

In order to calculate a confidence interval for this estimate of the correlation in direction of color evolution between each pairing of the four rivers, we first bootstrapped whole-fish color patterns. Populations were resampled 1000 times using the *boot* function in the R *boot* package (Davison and Hinkley 1997, Canty and Ripley 2019). Whole fish color patterns were bootstrapped in order to retain the associations of RGB values within and among the sampling points. For each bootstrap sample, we calculated the population mean color pattern as described above. This produced 1000 bootstrap estimates of the population mean color pattern. To generate 1000 estimates of the direction of evolution within a river, we subtracted the mean vector of the first bootstrap sample in the high-predation population from the mean vector of the first bootstrap sample in the low-predation population; this was repeated for the remaining 999 bootstrap samples from the high- and low-predation populations. In this way, we produced 1000 estimates of direction of color evolution for a given river. We then determined the similarity between the vectors of color evolution for each pair of rivers by calculating the vector correlation between each of the 1000 bootstrap estimates of color evolution within the rivers. Pair-wise comparisons of four rivers generated six distributions of correlation coefficients.

To determine whether the correlation between pairs of rivers was more similar than expected by chance, we calculated an expectation for the correlation between random vectors. We first generated vectors to simulate two rivers where the expected direction of evolution within each river was random. This was done using the *mvrnorm* function in the *MASS* package in R (Venables and Ripley 2002). We created two matrices, each with 1000 random vectors of length = 7386, mu=0 (vector giving the mean of zero for each variable of the matrix), and Sigma defined as the identity matrix (ones on the diagonal and zeros on the off-diagonal) to simulate random direction of evolution estimates. Similar to our bootstrap samples, we then paired each vector between the two simulated river estimates and calculated the mean correlation between random vectors. If the estimate of the vector correlations for each river comparison was above or below the mean correlation between random vectors, we determined the correlation between rivers in the direction of color evolution was more correlated than expected by chance.

### Phenotypic variance in color between predation regimes

We compared phenotypic variation in male coloration between high- and low-predation populations using the same rivers included in our analysis of the direction of color evolution (Aripo, El Cedro, Guanapo, and Turure). Using the data where the RGB values were collected using the 4DT, sampling circle diameter = 1 pixel sampling scheme (described above), we calculated the trace of the variance-covariance matrix to determine the phenotypic variance in color for each real population.

We used permutation to test if the phenotypic variation differed between high- and low-predation populations within a river. We permuted the label of high-predation or low-predation for each whole-fish color pattern within a single river using the *sample* function in base R (V3.5.3), and created 1,000 permutation samples for each river in this manner. For each permuted sample, we calculated the variance-covariance matrix (as described above) for the high- and low-predation permuted samples and calculated the trace to reflect the color pattern variance. We then subtracted the trace of the high-predation population from the trace of the low-predation fish for each permuted sample. The distribution of these 1000 measures of difference in variance was used to represent the null distribution. We calculated the observed population differences in total phenotypic variance from our real data and tested this against our null distribution. Using a two-tailed distribution, we calculated the proportion of values in the null distribution that were greater than, or less than, the observed difference in color pattern variance between the high- and low-predation populations to indicate the significance of the difference.

We were also interested in characterizing the spatial pattern of within-population variance in color. To do so, we compared the within- and between-population components of variance for each color channel at every x,y coordinate in the data set. We estimated variance components with restricted maximum-likelihood using *Proc Varcomp* of SAS version 9.4 (SAS Institute, Cary, NC), with the color value (R, G, or B) at a given x,y position as the dependent variable, year as a fixed effect, and population ID as a random effect.

## Results

### Multivariate differentiation among populations

We first sought to determine a combination of sampling density and sampling circle size that would enable us to classify the 11 guppy populations with the highest precision. To do so, we compared the cross-validation results for the 12 sampling designs (combinations of three sampling schemes and four sampling circles sizes). Here, we report the results from analyses performed when images taken in different years were separated (see Supplemental Table S3 for results from analyses when images from both years were combined). As the number of rounds of Delaunay triangulations (DT) increased, the RMSE, averaged over all sampling circle sizes, consistently decreased (RMSE averaged across years: 2DT = 0.18, 3DT = 0.15, 4DT = 0.13; Supplemental Table S3), and the mean successful assignment of the validation set increased (proportion of successful assignment averaged across years: 2DT = 0.87, 3DT = 0.90, 4DT = 0.91; Supplemental Table S3). To visualize the difference in sample density, we reproduced an example image by plotting the sampled RGB values at each of the three sampling densities (Figure 2C); clearly, the increased sampling density improved the representation of the complex guppy color pattern. When images were reproduced using color sampled at the four different sampling circle diameters (1,3,5, and 9 pixels), there was no noticeable difference in image quality that would suggest that one of the four sampling circle diameter sizes would best represent an image (Supplemental Figure S3). In order to determine our sampling circle diameter for our analyses, we evaluated the cross-validation results when images were combined and when results were averaged across years. At the highest sampling density (4DT), classification success was high and nearly constant irrespective of sampling circle diameter (range: 0.91 – 0.93). We therefore decided to use sampling circle diameter of 1 (1 pixel sampled, no pixel averaging) because it produced the lowest RMSE and highest success rate in both years, when years were analyzed separately (Supplemental Table S3). This sampling scheme produced a color vector of length 7386 (2462 sampling points * 3 color channels).

Using this sampling scheme, DAPC analysis generated discriminant functions representing principal components that best differentiated guppy populations. Figure 3 shows the coordinates of all individual fish within the 11 populations plotted on the first three discriminant function axes. Discriminant function axes 1, 2, and 3 accounted for 45.6%, 13.6%, and 13.2% of the variation in color patterns, respectively (Figure 3). Axis 1 separated populations predominantly by the year in which they were sampled; populations collected in 2016 were clustered to the left and did not overlap with the samples collected in 2017, which were clustered to the right in the scatterplot (Figure 3A). In contrast, discriminant function axis 2 consistently separated high- and low-predation populations of the same river, with high-predation populations higher (more positive) on axis 2 than the corresponding low-predation population (Figure 3A). Indeed, even though the two Marianne populations were sampled in different years and there were slight differences in the resolution of the images, the direction of separation along axis 2 followed this pattern. Discriminant function axis 3 also consistently separated high- and low-predation population pairs, with high-predation populations always placed higher (more positive) on axis 3 than their low-predation counterparts (Figure 3B). In Figure 3B, which plots axes 2 and 3, it is evident that these two axes together consistently differentiate low- and high-predation population pairs, with low-predation populations always below and to the left (more negative on both axes) of their high-predation counterpart.

**Figure 3.**
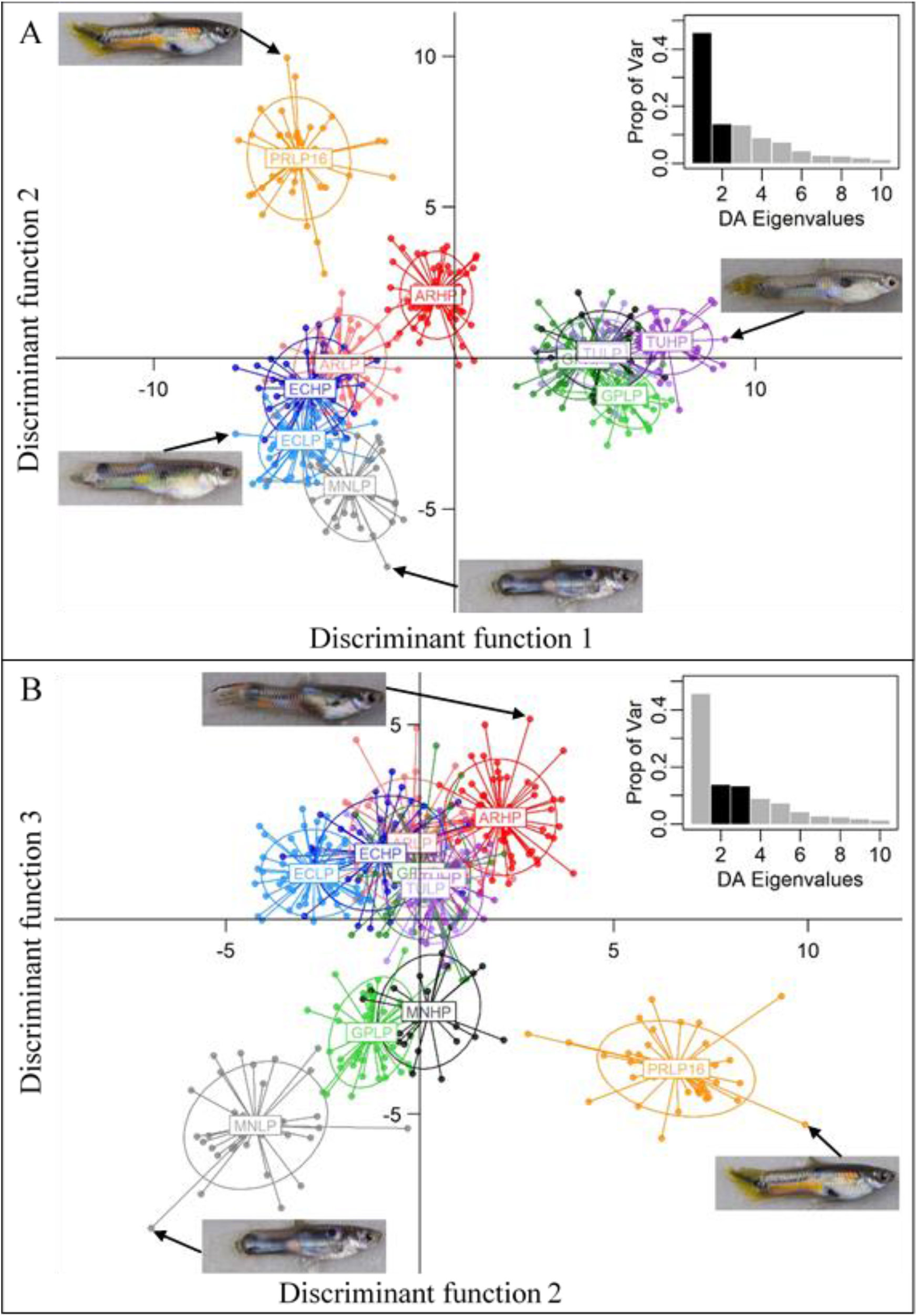
Scatter plot from the discriminant analysis of principal components (DAPC) where population membership was predefined (11 populations). Points represent the coordinates of individuals and populations are within inertial ellipses. The four letters at the center of the inertial ellipses represent the population: the first two letters identify the river (AR=Aripo, EC = El Cedro, GP = Guanapo, MN = Marianne, PR = Paria, TU = Turure), followed by two letters that identify the predation regime (HP = high-predation, LP = low-predation). High- and low-predation populations sampled within the same river are represented by dark and light versions of the same color, respectively. The barplot (inset) shows the proportion of variation explained by the discriminant analysis eigenvalues; dark bars correspond to the axes presented in the scatter plot of each panel section. Discriminant function axes one and two are plotted in panel (A) and axes two and three are plotted in panel (B). Original images (prior to unwarping) of individuals located at the extremes of each axis are shown for each panel. The color data used for the DAPC was collected from photos unwarped to a consensus shape and red, green, and blue (RGB) color channels were sampled from 2462 pixels.

Because axes two and three presented a consistent pattern of separation associated with predation regime differences within river, we were interested in visualizing the locations of the color samples that contributed most to the separation along these axes. For each axis, the top 10% of the color values (N=739) that contributed to each of these discriminant function axes are plotted within the outline of the consensus fish shape (Figure 3). However, each fish plot shows fewer than 739 points since sampling locations may be represented by more than one color channel (R, G, and B). Sampling locations that contributed most to separating populations along axis 2 were clustered on the caudal peduncle area and mid body anterior to the dorsal fin, while those along axis three included some areas of the caudal peduncle, mid body, and locations around the eye of the fish. Very few sample points on the caudal fin contributed to discrimination among populations on either axis.

### Comparison of direction of evolution within and between rivers

The natural replication of the ancestor-descendant relationship between rivers allows us to determine whether the direction of color pattern evolution is more similar between rivers than expected by chance. The DAPC analysis described above suggests that the direction of evolution is somewhat consistent across drainages, but we can test this hypothesis directly by examining the direction of changes in the full data space. To do this, we tested whether the direction of color pattern evolution is more consistent across different rivers than expected by chance.

Figure 4 illustrates the similarity in direction of color evolution among the four rivers by showing the pairwise correlation between vectors that describe the average color pattern evolution within each river. Each pair of rivers (e.g., Aripo vs El Cedro) is represented by a bootstrap distribution that shows the estimate of the correlation and the uncertainty in that estimate in the direction of evolution between rivers (95% Confidence Interval). Of the six pairwise comparisons, the El Cedro-Guanapo (observed = 0.331, CI = 0.150 to 0.381) and the Aripo-Turure (observed = 0.376, CI = 0.133 to 0.418) pairs were more correlated than expected by chance (mean correlation of random vectors = 0.002; Figure 4). The remaining four pair-wise comparisons were not more correlated than expected by chance: Aripo-El Cedro (observed = 0.105, CI = −0.102 to 0.280), Aripo-Guanapo (observed = 0.062, CI = −0.070 to 0.163), El Cedro-Turure (observed = −0.094, CI = −0.259 to 0.132), Guanapo-Turure (observed = 0.006, CI = −0.127 to 0.131).

**Figure 4.**
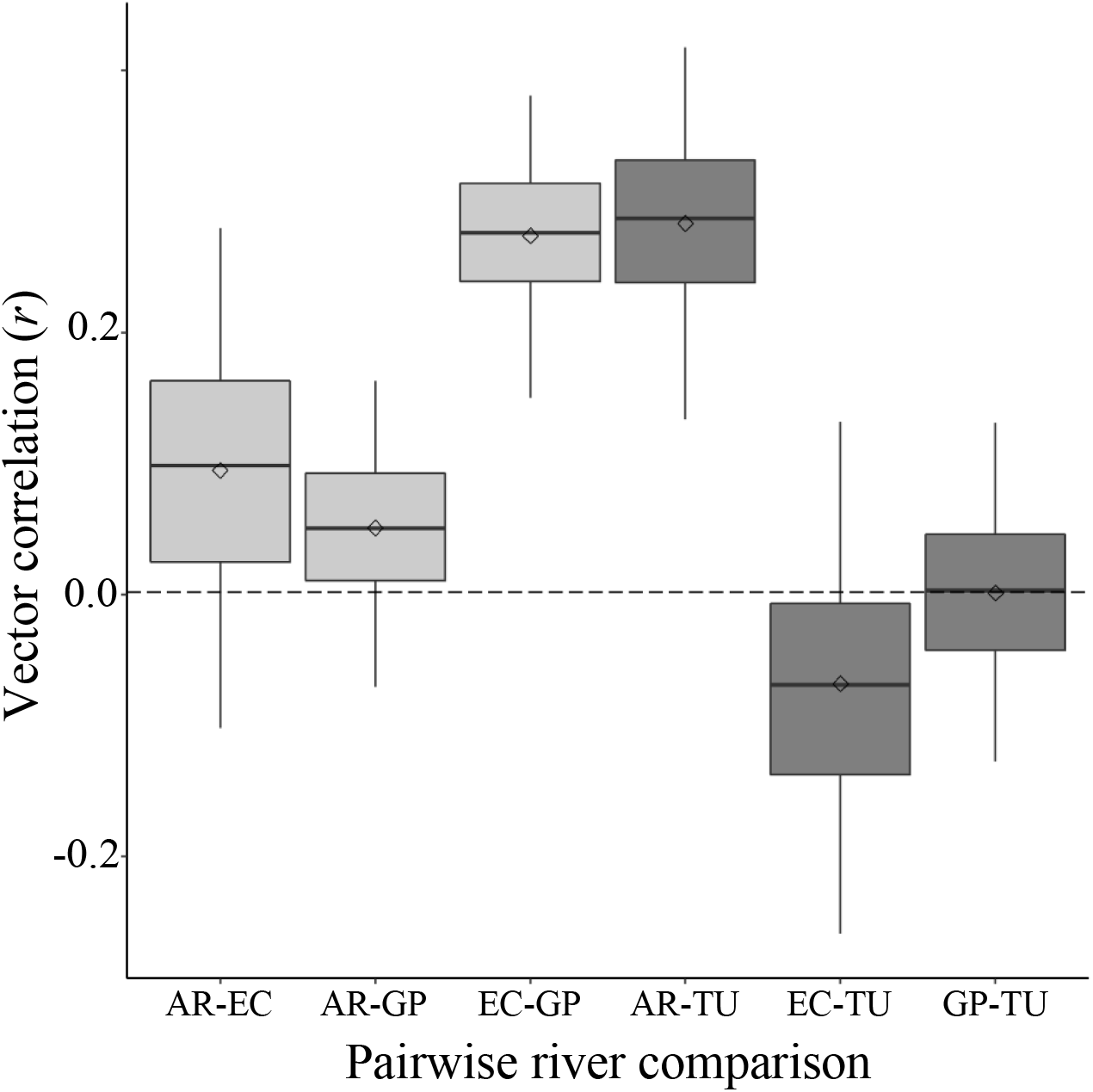
Correlation in direction of evolution between pairs of rivers. Pair-wise river comparisons are represented by each box and whisker where river pairs are identified on the x axis. Boxes indicate the interquartile range and whiskers show the 95% confidence interval of the estimate for the correlation between vectors of direction of color evolution. Within each boxplot, the median and mean of the distributions are represented by the solid line and diamond, respectively. The horizontal dashed line represents the mean correlation between random vectors. The horizontal dotted line represents the mean expected correlation between pairs of rivers within the Caroni drainage system. The four rivers included in the pair-wise comparisons are the Aripo (AR), El Cedro (EC), Guanapo (GP), and Turure (TU). AR, EC, and GP belong to the Caroni drainage and TU to the Oropuche; within drainage pair-wise comparisons are shaded in light grey while between drainage comparisons are shaded in dark grey.

Of the four rivers included in our analysis of direction of color evolution, three rivers belonged to the Caroni (Aripo, El Cedro, and Guanapo) and one the Oropuche (Turure) drainage. This allowed for a comparison of whether correlations in direction of color evolution between rivers belonging to different drainages were weaker than correlations within drainage. The cross drainage correlations were somewhat more variable than the within drainage correlations; interestingly, the Aripo-Turure correlation is just as strong as the strongest within drainage correlation (Figure 4).

### Comparison of phenotypic variance in color between predation regimes

We next compared the total phenotypic variation between predation regimes for the four rivers in which both regimes were sampled in the same year, using the 4DT, 1 pixel diameter sampling scheme. Figure 5 shows the results of the permutation tests of the difference in variance between high and low predation populations within a river. In two rivers (Aripo and Turure), the low-predation population had significantly higher total phenotypic variance (permutation P-value < 0.001 and P=0.009, respectively). In the other two rivers (El Cedro and Guanapo) the high-predation population had nominally higher total phenotypic variance, but these differences were not significant (permutation P-value = 0.099 and P=0.113, respectively; Figure 5).

**Figure 5.**
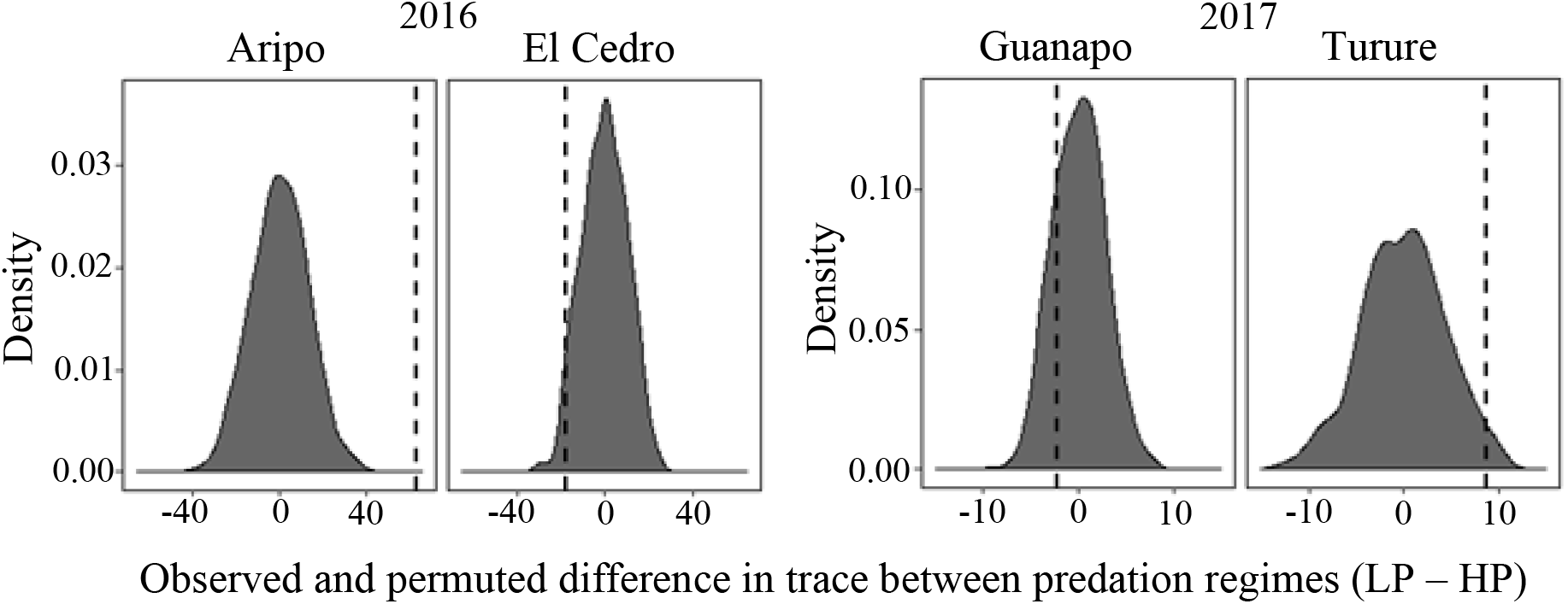
Observed and permuted difference in total phenotypic variance (trace) between predation regimes where high-predation (HP) values were subtracted from low-predation (LP) values within a river. Observed difference in trace is given by the vertical dashed line on the same plot as the distribution of the permuted differences in trace between predation regimes for a river.

A spatially-explicit analysis revealed the positions on the body that varied most within populations, relative to among-population variance (Figure 6). For example, a position dorsal to the pectoral fin (“shoulder spot”), and several positions in the posterior region of the caudal peduncle were highly variable among populations. Conversely, in a region immediately posterior to the “shoulder spot”, most variation was distributed between, rather than within populations, as indicated by low values in the heat maps in Figure 6. Notably, these regions did not differ substantially in overall variance, suggesting that these patterns were driven by the distribution of variance within and among populations, and not by differences among body regions in general variability (Supplemental Figure S4).

**Figure 6.**
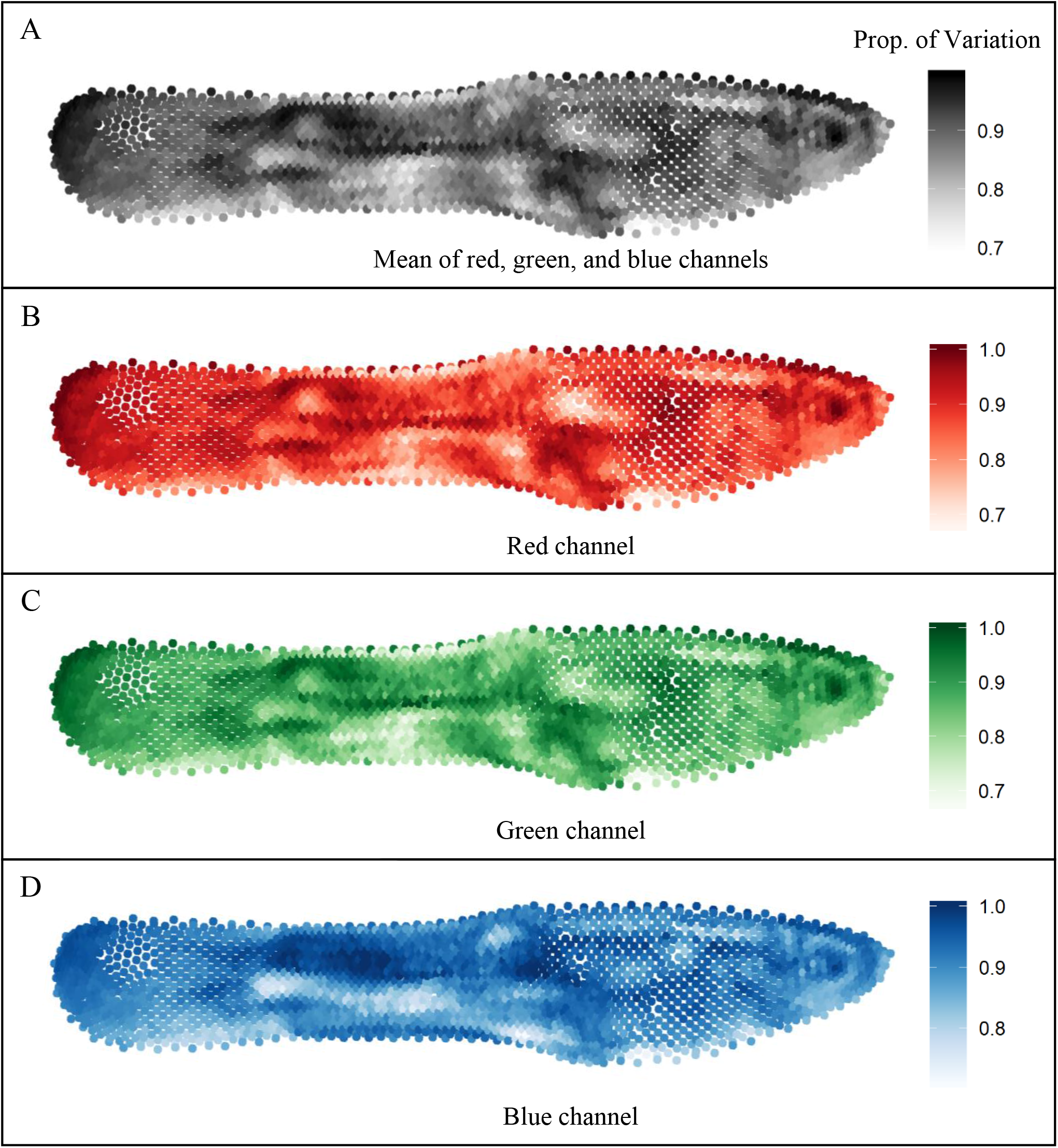
A heat map showing the proportion of variation accounted for by differences within each of the 11 populations in color sampled at each of the 2462 sampling points. Darker colors indicate higher proportion of variance distributed within, rather than among popuations. Panel A shows the mean of the proportion of variation given by the red (B), green (C), and blue(D) channels.

## Discussion

Our approach to quantifying variation in color patterns addresses important limitations in the tools currently available for color pattern analysis. Colormesh allows the analysis of variation in chromatically and spatially complex patterns, avoids loss of information by catagorizing color or identifying color elements, allows high-throughput analysis by using digital images of whole color patterns, and provides flexiblity with respect to sampling density and area. We used this novel method to determine if it could accurately assign fish to the correct populations and to evaluate whether the direction of color pattern evolution was consistent in four different rivers. We also asked if total phenotypic variation differed consistently between up- and down-stream populations across drainages. Below we discuss the performance of our approach on these tasks and compare our conclusions to those arrived at by using different methods.

### Population differentiation

Our approach allows the color of an individual to be characterized at different numbers of points on the body, and for different numbers of pixels around each point. Therefore, an important precursor to analysis is to investigate how varying these two parameters affects the conclusions of any analysis. For our guppy samples, increasing the number of sampled points increased our ability to differentiate populations, but varying the number of pixels averaged around each point made little difference. This is likely because the guppy color pattern is complex with color changing fairly rapidly and dramatically at small spatial scales. Organisms with less fine-grained patterns might be sufficiently sampled at courser scales.

DAPC using our selected sampling scheme (4DT, 1 pixel diameter) separated populations largely by sampling year along the first discriminant function axis. Camera settings differed between years which probably accounts for the variation explained by this axis; however, since different populations were sampled in different years, this axis probably also captured some population differences in color. Although this year effect limited some of the downstream analyses we could perform, it also indicated that this method is sensitive to unsuspected sources of variation, which can be a valuable tool in assessing data quality.

DAPC also allowed us to identify axes of variation that consistently separated high- and low-predation populations along discriminant function axes two and three, although the degree of differentation varied among river systems. Differences between predation regimes have been reported in previous studies assessing color patterns; however, the nature of these reported differences have varied. For example, in a study of 112 sites sampled from 53 streams located in Trinidad and Venezuela, Endler (1978) reported that males in high-predation natural populations had smaller and fewer spots, and less coverage of bright (iridescent) colors. In contrast, Millar and Hendry (2012) reported few consistent differences between high- and low-predation populations across five different river systems even though methods of color quantification (color categorization and measures of discrete color spots as judged by human vision) were similar to those used by Endler (1978). Investigations of color evolution across a similar ecological gradient, but over short time scales, have also produced diverse results. Using populations translocated from high-predaton to low-predation locales, and a combination of spectrometry, digital photography, and visual modeling, Kemp and colleagues reported that color evolved along different trajectories in the Aripo and El Cedro river introductions (Kemp et al. 2008, 2009). While there were differences in color quantification methods in these studies, strong ecological correlates of color divergence suggest that these differences between river systems are driven by biology, not methodology (Millar et al. 2008; Millar and Hendry 2012; Kemp et al. 2018). Our analysis of the multivariate direction of evolution (see below) provides a quantitative measure of degree of similarity or parallelism in independent replicates using methods developed for addressing these questions and similar questions for other kinds of quantitative traits.

### Direction of color evolution

We sought out to test whether the repeated transition between high- to low predation pressure among rivers resulted in parallel directions of color pattern evolution among several rivers. Using the known ancestor-descendant relationship between high- and low-predation populations, we calculated correlations in direction of multivariate color evolution between rivers. Results using our novel approach to whole color pattern analysis, were similar to those reported in two previous studies that found differences among the rivers included in our study in the direction of color evolution; in both studies, directional difference in quantity and size of individual color pattern elements were compared to determine similarity in evolutionary trajectories (Kemp et al. 2009, Millar and Hendry 2012). Three of the four rivers (Aripo, El Cedro, and Guanapo) in our comparison belong to the same drainage system (Caroni) and the El Cedro is a tributary of the Guanapo. Comparisons between rivers within the Caroni drainage system found that the El Cedro and Guanapo rivers showed more similarity in the trajectory of evolution than the other pairs of rivers, but the degree of similarity was only moderate (vector correlation <0.3).

Between drainage correlations included comparisons of a single river of the Oropuche drainage (Turure) with each river belonging to the Caroni drainage. These three correlation measures were highly variable with one comparison, the Turure-Aripo, having moderately similar trajectories in direction of color evolution. The low-predation Turure population was originally a guppy-free location where guppies from a high-predation population in the Arima river (in the Caroni drainage, collected near the confluence with the Guanapo river) were introduced by Haskins in 1957 (Magurran et al. 1992, Shaw et al. 1992). Genetic evidence indicates the introduced guppies have successfully established themselves along the low- to high-predation reaches of the Turure and have interbred with native Turure populations located downstream (Shaw et al. 1992, Becher and Magurran 2000, Fitzpatrick et al. 2015). We therefore might have expected the Turure to show a similar direction of evolution to the the Guanapo and the El Cedro. However, the between drainage system comparison found the direction of color evolution between the Turure and Aripo was more similar to that expected between rivers within the same drainage; as expected, the Turure-Guanapo and Turure-El Cedro correlations were weaker.

In our study, the consistent pattern of separation between high and low-predation poulations found in the DAPC suggested that the direction of color evolution might be quite similar in different rivers. It is important to note that our approach sampled all aspects of color data obtained by the digital imaging process. It is possible that evolutionary trajectories of some aspect of color are indeed more parallel but remain undetected. However, our method does provide the user with the option of restricting analysis to one color channel, or to a single aspect of a different colorspace such as HSL (Hue, Saturation, and Lightness), to test for parallel evolution among specific aspects of color. The ability to quantify parallelism in this way thus allows more nuanced view of parallelism.

### Total phenotypic variation

Finally, we evaluated differences in total phenotypic variation within and between predation regimes. Male guppy color patterns are among the most genetically variable phenotypes reported in the literature (Brooks and Endler 2001b, Hughes et al. 2005), and the processes that promote variation have been investigated extensively (e.g., Farr 1977, Houde and Endler 1990, Hughes et al. 1999, 2013, Brooks and Endler 2001, Brooks 2002). Results from field and laboratory experiments indicate that both predators and females exert negative frequency dependent selection (NFDS) on male color patterns, which promotes high phenotypic and genetic variation in color (Hughes et al. 1999, 2013, Brooks and Endler 2001a, Olendorf et al. 2006, Fraser et al. 2013). *Rivulus hartii*, which preys on male guppies and which is more common in low-predation sites, appear to learn to be more effective guppy predators the more they are exposed to a particular color pattern (Fraser et al. 2013). In natural populations, males bearing rare color patterns have higher survival than males bearing common patterns, and this effect is stronger in low-predaton sites that have many *R. hartii* (Olendorf et al. 2006). Female guppies also prefer males bearing rare or unfamiliar color patterns (Eakley and Houde 2004, Zajitschek et al. 2006, Zajitschek and Brooks 2008, Hampton et al. 2009, Mariette et al. 2010, Hughes et al. 2013, Graber et al. 2015), and this preference has been documented in both high- and low-predation sites (Valvo et al. 2019). Taken together, these results suggest that low-predation sites should exhibit more variation in male color patterns, since NFDS by *R. hartii* should be stronger, and sexual selection by female preference is equally strong in the low- and high-predation sites. To our knowledge, there has previously been no attempt to quantify whole-color pattern variation that would allow this prediction to be tested.

We found that the low-predation Aripo and Turure populations had significantly greater variance in overall color than the high-predation populations in those rivers. Although not significant, the trend was in the opposite direction for the El Cedro and Guanapo populations. The lack of consistently greater variation in low-predation populations suggests that *R. hartii* might not inflict strong NFDS on guppies in some natural populations, or that other ecological conditions interact with the NFDS selection imposed by predators and females to determine levels of within-population variance in male color. Future investigations should assess density of potential predators and other ecological and genetic patterns (e.g., effective population size) to provide more insight into the determinants of variation in this ecologically important trait.

We also identified spatial positions on the body that were characterized by high within-population variation. Both natural and sexual selection favor males with rare or unfamiliar color patterns in this species (Hughes 1999; Zajitschek and Brooks 2008; Olendorf et al. 2006; Hughes et al. 2013), imposing negative frequency dependent selection. Future studies could examine whether spatial positions exhibiting high within-population variance are subject to stronger selection than regions with lower variance. Such studies could determine if females-imposed sexual selection, (e.g., Valvo et al. 2019) or predator-imposed natural selection (e.g., Fraser et al. 2013) are influenced by the spatial distribution of variable color pattern elements.

### Summary, extensions, and limitations

The novel method of color analysis described here adds to the toolbox of methods for color pattern quantification. In applying this pipeline, we focused on population differences and on patterns associated with repeated evolutionary transitions to novel ecological conditions in guppies. However, this method of quantifying color, which results in a standard multidimensional respresentation, can be used to address a wide variety of questions, and can be integrated or combined with other approaches to color and pattern analysis. For example, patterns of correlation in color across the body could be assessed by standard methods in spatial statistics (Schabenberger and Gotway 2005, Wackernagel 2013). These methods can identify regular patterns such as stripes and spots, and capture variation in this patterning among individuals, populations, or species. Our approach can be easily extended to more than three color channels (e.g., UV), to different color spaces (CIELAB, HSV, etc.), and to other features such as polarization by adding dimensions to the multivariate trait vector. The continuous representation of color could be used for image segmentation or disparity analysis, using approaches available in *patternize* and *colordistance* (Van Belleghem et al. 2018, Weller and Westneat 2019). Critically, the highly multivariate and spatially-explicit data produced by this method could be integrated with visual models based on the biology of the receivers of visual signals, such as those incorporated in *pavo2* and the QCPA platform (Maia et al. 2019, van den Berg et al. 2020). Visual modeling could resolve the apparent contrast between our quantitative results (low to moderate parallelism in the evolution of color patterns across different river systems) with the results of other studies that find that rivers within the same drainage tend to exhibit parallel changes (e.g., Kemp et al. 2019). One possible cause of this disparity is that color elements that are perceived by predators and mates tend to evolve in parallel, while other elements do not.

This method of color pattern analysis may also be used to investigate the inheritance of the complex guppy color pattern. Some of the genes responsible for coloration are located on the sex chromosomes with several of these genes being Y-linked (Winge 1927, Lindholm and Breden 2002). The guppy color pattern is often discribed as being so variable, that no two fish look alike; however, some aspects of the color pattern are highly heritable (Brooks and Endler 2001b, Hughes et al. 2005) and some appear to be inherited patralineally (Winge 1927, Endler 1978, Houde 1997). The extent to which guppy color patterns are sex-linked (as opposed to sex limited in expression), and how these patterns relate to chromosomal evolution, have been the subject of considerable recent interest (Gordon et al. 2012, 2017, Wright et al. 2017, Charlesworth 2018). The methods described here could be deployed to quantify the number of different color patterns that segregate within different natural populations and, when combined with appropriate breeding designs, could distinguish color pattern components that display autosomal and sex-linked patterns of inheritance.

Our method is based on unsupervised color quantification combined with geometric morphometrics to identify regions of putative homology across individuals and populations. The combination of flexibility to accommodate different sampling schemes, along with completely objective sampling allowed us to address novel questions about the evolution of complext color patterns in guppies, such as comparing evolutionary trajectories and comparing phenotypic variance at a highly multivariate scale.While powerful and flexible, this approach does have limitations. For example, substantial shape difference between specimens could decrease the accuracy of the homology alignment. Finally, several authors have noted that assessment of structural coloration is challenging (Endler 1990, Kemp et al. 2008, 2018, Vukusic and Stavenga 2009). While we did not address this issue here, we note that, like the addition of UV or polarization information, incorporating meaures taken under different lighting or viewing condtions would require no fundamental change to the approach described here. These additional features will increase the dimensionality of the data and therefore the computing resources required. However, all calculations reported here were conducted on a modestly powered desktop PC.

## Acknowledgements

This work was supported by a Rosemary Grant Award (2015) to J.J.V., two National Science Foundation Grants (DEB 1740466 and OIS1354775) to K.A.H., and funding by the Natural Sciences and Engineering Research Council (Canada) to F.H.R.

We would like to thank Connor Fitzpatrick, Michael Foisy, Alex De Serrano, Mark Charran, Jack Torresdal, Diana, and Tuna-puna for their assistance doing the field collections. We are grateful to Ronnie Hernandez at the William Beebe Tropical Research Station (Trinidad) for his assistance in providing supplies and accommodations required for completion of fieldwork over the two-year period, and we thank the government of Trinidad and Tobago for permitting the collection of experimental populations. We also thank Alexa Guerrera and Amber Makowicz for their helpful comments that substantially increased the quality of this manuscript.

