## Supplemental_material for "Colormesh: A novel method for quantifying variation in complex color patterns"

### Methods

#### *Fish collection*

Fish were caught using butterfly nets and transported in sealed Nalgene bottles containing water with Stress Coat (API) to the nearby William Beebe Tropical Research Station, located in the lower Arima valley in the Northern Range, Trinidad. Fish were separated by population and maintained in single-sex 20-40L aquaria for 24-48 hours prior to taking their photographs. Water exchanges of 30% were performed daily using conditioned rainwater collected on site. Fish were fed Tetramin Tropical flake food twice daily.

#### *Image collection*

To compare color patterns within and among rivers, we collected digital photographs of male guppies by lightly anesthetizing them with buffered MS222. Males were placed on their side, with the snout facing right, on a stage along with a scale and color standard. Using a camel-hair paintbrush, we positioned the dorsal fin and gonopodium away from the body of each fish and dabbed the fish body with a Kimwipe to remove excess water and reduce glare. They were photographed (Canon EOS 5D, 100mm f/2.8 macro lens) within a Photo Cube lighting tent to diffuse light and minimize glare. Three 65-watt daylight (white) compact fluorescent light bulbs (Fovitec StudioPRO, 5500K, full spectrum, Color Rendering Index = 90) in hooded light fixtures were positioned outside of the lighting tent. Two were positioned opposite each other, to the left and right of the photographer and angled down approximately 30 degrees from the plane of the camera lens (see Supplemental Figure S1 for photography setup); the third light source was positioned on the side of the tent opposite from the photographer and angled down approximately 20 degrees. The configuration of the three lights minimized any shadow. Images were taken in RAW format and converted to TIFFs in Adobe Photoshop CC 2018. A minimum of two photos of each fish were collected and photographs of all males from the same population were taken during the same session. We photographed N=285 male guppies in 2016 and N=200 in 2017 for a total of N=485 images. All fish were photographs on an 18% grey background (Movo Photo Color/White Balance Card Set for digital photography). The photography equipment used in 2016 and 2017 was identical. However, the distance of the camera lens from the fish differed between years. We increased the distance between the stage and camera from 14.5cm in 2016 to 28cm in 2017 which allowed for the use of autofocus and a remote switch for a more consistent image quality. Because of this difference, we took year into account in subsequent analyses. For a sample of images from each of the 11 populations, see Supplemental Figure S2.

### Tables & Figures

Supplemental Table S1. River/tributary name(s) and predation regime, year in which males were collected, the total number of males sampled, river system drainage membership, and GPS coordinates of populations sampled.

| River/predation regime | Year | Total sampled | Drainage | Latitude – N | Longitude - W |
|---|---|---|---|---|---|
| Aripo high-predation | 2016 | 57 | Caroni | 10.65474 | 61.22755 |
| Aripo low-predation (main river) | 2016 | 51 | | 10.67123 | 61.22922 |
| El Cedro high-predation | 2016 | 47 | Caroni | 10.6567 | 61.26599 |
| El Cedro low-predation (Experimental) | 2016 | 54 | | 10.663588 | 61.26584 |
| Guanapo high-predation (Twin Bridges) | 2017 | 57 | Caroni | 10.63989 | 61.24833 |
| Guanapo low-predation (Tumbasson) | 2017 | 45 | | 10.70944 | 61.25778 |
| Marianne high-predation | 2017 | 24 | Northern | 10.76667 | 61.30000 |
| Marianne low-predation | 2016 | 36 | | 10.75727 | 61.31523 |
| Paria low-predation | 2016 | 40 | Northern | 10.74740 | 61.26629 |
| Turure high-predation (Experimental) | 2017 | 40 | Oropuche | 10.65469 | 61.16946 |
| Turure low-predation (Experimental) | 2017 | 34 | | 10.68606 | 61.17312 |

Supplemental Table S2. Name, color code and RGB values for the Liquitex Heavy Body acrylic paint used for the color standards. RGB values for pixel colors range between 0 and 1, therefore RGB values were divided by 255 to determine the value to correct the measured values at each sampling point.

| Liquitex Color Name | Code | Red | Green | Blue |
|---|---|---|---|---|
| Napthol crimson | 292 | 173/255 | 43/255 | 50/255 |
| Cadmium orange | 150 | 243/255 | 121/255 | 33/255 |
| Cadmium yellow medium | 161 | 254/255 | 244/255 | 17/255 |
| Emerald green | 650 | 43/255 | 163/255 | 73/255 |
| Ivory black | 244 | 26/255 | 13/255 | 21/255 |

Supplemental Table S3. Below are the results from the *xvalDapc* function in the *adegenet* package using the number of PC's associated with the lowest Root Mean Squared Error (RMSE) for each of the 12 different sampling schemes. Also given is the proportion of successful placement of the validation set and the number of PC's retained. The sampling designs evaluated include all combinations of three sampling densities (2, 3, and 4 rounds of Delaunay triangulations (DT)) and four different sample circle sizes (diameter in pixels). RGB values of all pixels within sample circles having a diameter > 1 were average to provide a single R, G, and B value. Cross validations were performed on datasets where images from both years were combined as well as separate.

| Year | Sample circle diameter (pixels) | 4DT | | | 3DT | | | 2DT | | |
|---|---|---|---|---|---|---|---|---|---|---|
| | | Success | RMSE | PC's | Success | RMSE | PC's | Success | RMSE | PC's |
| 2016 & 2017 | 1 | 0.913 | 0.109 | 100 | 0.900 | 0.120 | 100 | 0.852 | 0.166 | 100 |
| | 3 | 0.924 | 0.098 | 150 | 0.917 | 0.101 | 100 | 0.861 | 0.160 | 100 |
| | 5 | 0.927 | 0.097 | 100 | 0.900 | 0.114 | 200 | 0.870 | 0.150 | 100 |
| | 9 | 0.919 | 0.101 | 150 | 0.913 | 0.102 | 150 | 0.881 | 0.135 | 150 |
| 2016 | 1 | 0.919 | 0.131 | 60 | 0.920 | 0.128 | 40 | 0.893 | 0.159 | 100 |
| | 3 | 0.912 | 0.132 | 80 | 0.894 | 0.153 | 60 | 0.910 | 0.150 | 60 |
| | 5 | 0.913 | 0.131 | 60 | 0.908 | 0.142 | 40 | 0.908 | 0.145 | 80 |
| | 9 | 0.908 | 0.138 | 60 | 0.918 | 0.125 | 80 | 0.902 | 0.140 | 120 |
| 2017 | 1 | 0.928 | 0.114 | 40 | 0.897 | 0.154 | 40 | 0.826 | 0.224 | 60 |
| | 3 | 0.900 | 0.142 | 60 | 0.892 | 0.153 | 60 | 0.808 | 0.232 | 20 |
| | 5 | 0.909 | 0.147 | 100 | 0.882 | 0.167 | 80 | 0.819 | 0.226 | 60 |
| | 9 | 0.925 | 0.121 | 80 | 0.894 | 0.154 | 80 | 0.862 | 0.195 | 40 |

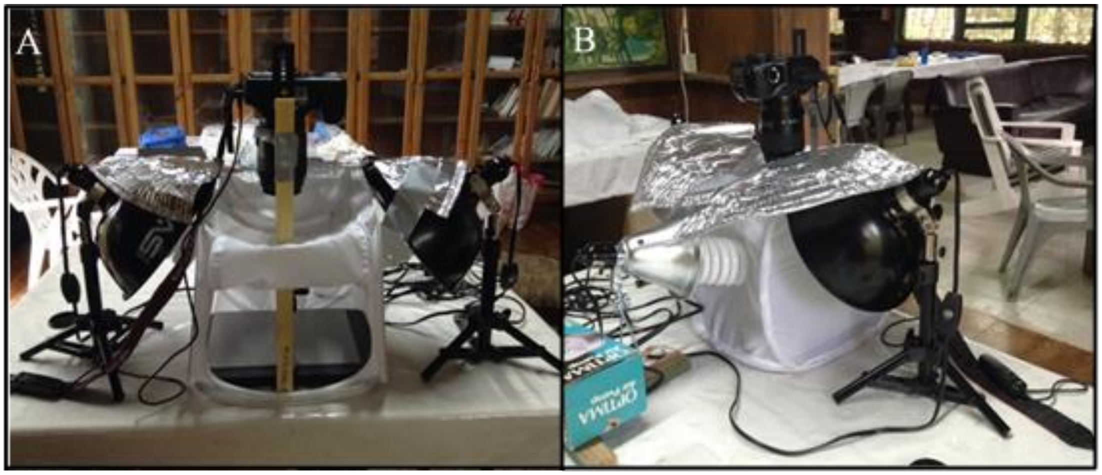


Supplemental Figure S1. Photography setup for digital image collection. Panels show the location of the three light sources. Individual fish were placed within the white photo tent through the opening visible in panel (A). Panel (B) shows the orientation of the third light source that is not visible in panel (A). The photography setup remained stationary for all sessions within a given year.

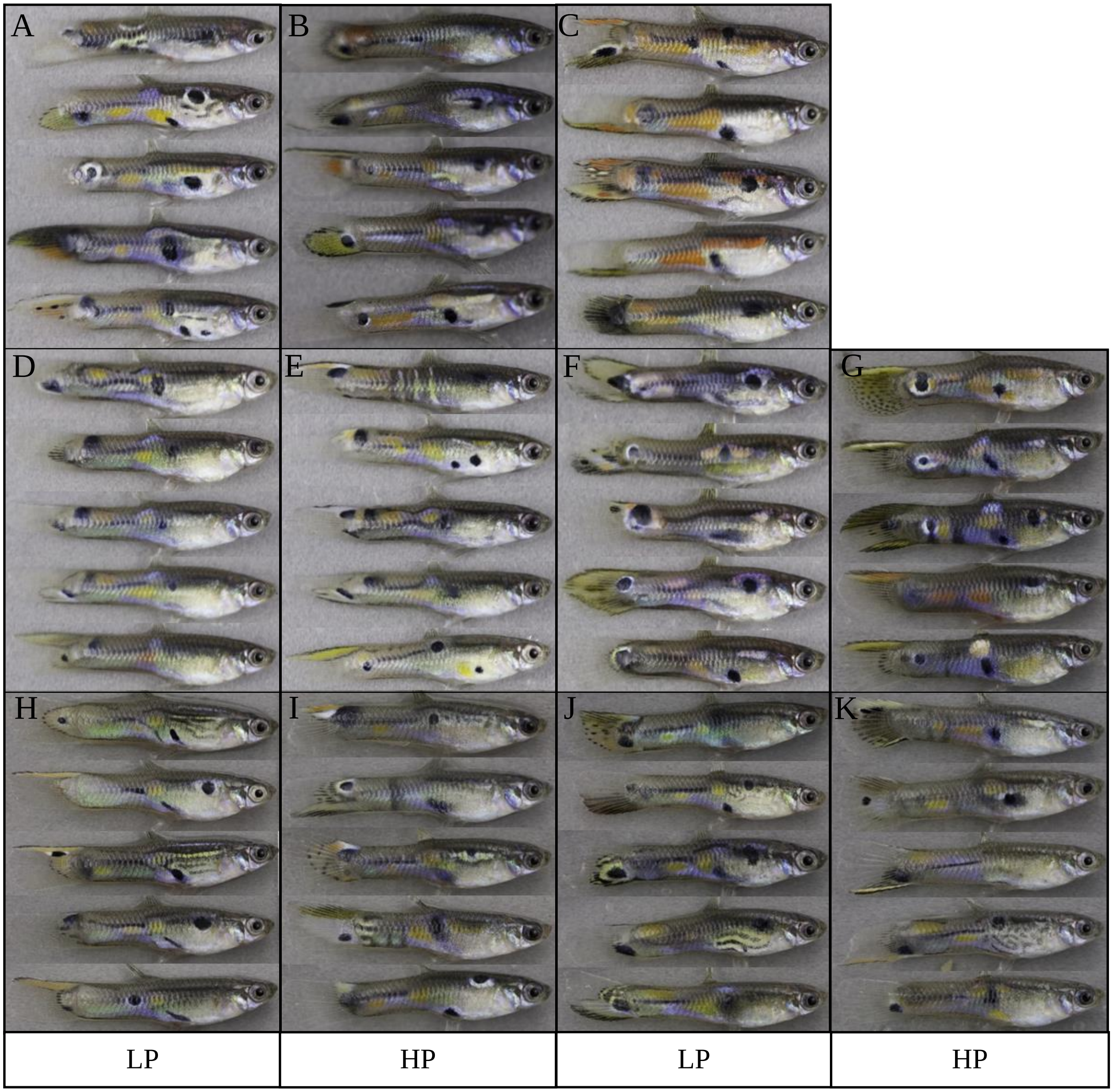


Supplemental Figure S2. Example images of six fish from each of the 11 populations sampled. Panels (A-F) were photographed in 2016 and Panels (G-K) were photographed in 2017. Populations are from the Aripo (A & B), Paria (C), El Cedro (D&E), Marianne (F&G), Guanapo (H&I) and Turure (J&K). Low predation (LP) populations are pictured to the left of high-predation (HP) populations in each row with the exception of Panel (C) since the Paria river lacks a high-predation contrast.

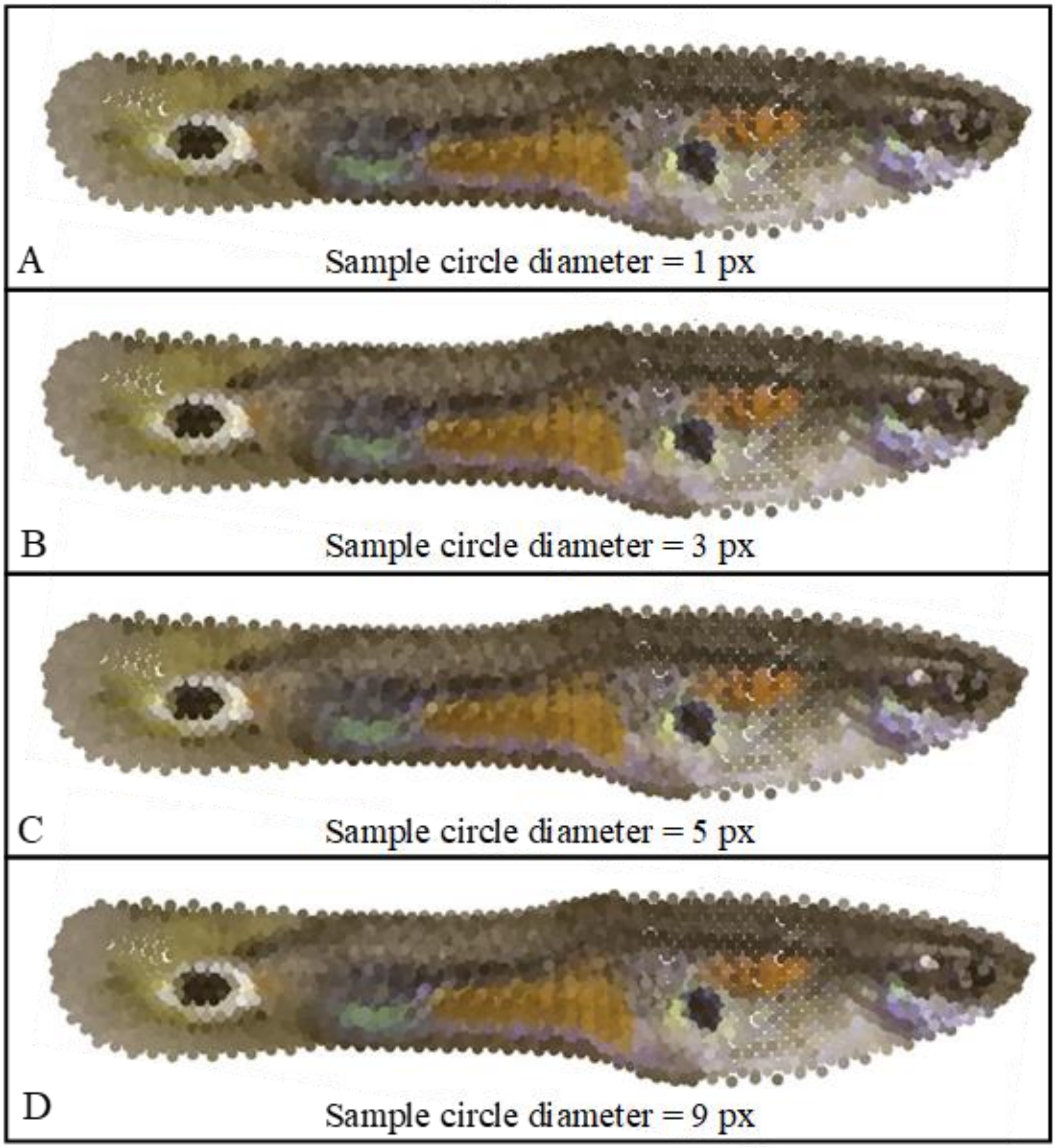


Supplemental Figure S3. Panels A – D show RGB color sampled using the four different sampling circle diameters. The density of sampling points in each panel was determined using four Delaunay triangulations. Panel (A) reproduced the RGB values samples from the 1 pixel (px) located at the centroid of each triangle. Panels B-D display the color produced from the mean red, green, and blue values of each pixel within the sampling circle of diameter = (B) 3 pixels, (C) 5 pixels, and (D) 9 pixels. For each of the four different sampling circle sizes, RGB values sampled at each location were plotted using the *plot* function in R. Plotted point size was equal (cex = 1.2) and selected to minimize white space between plotted points.

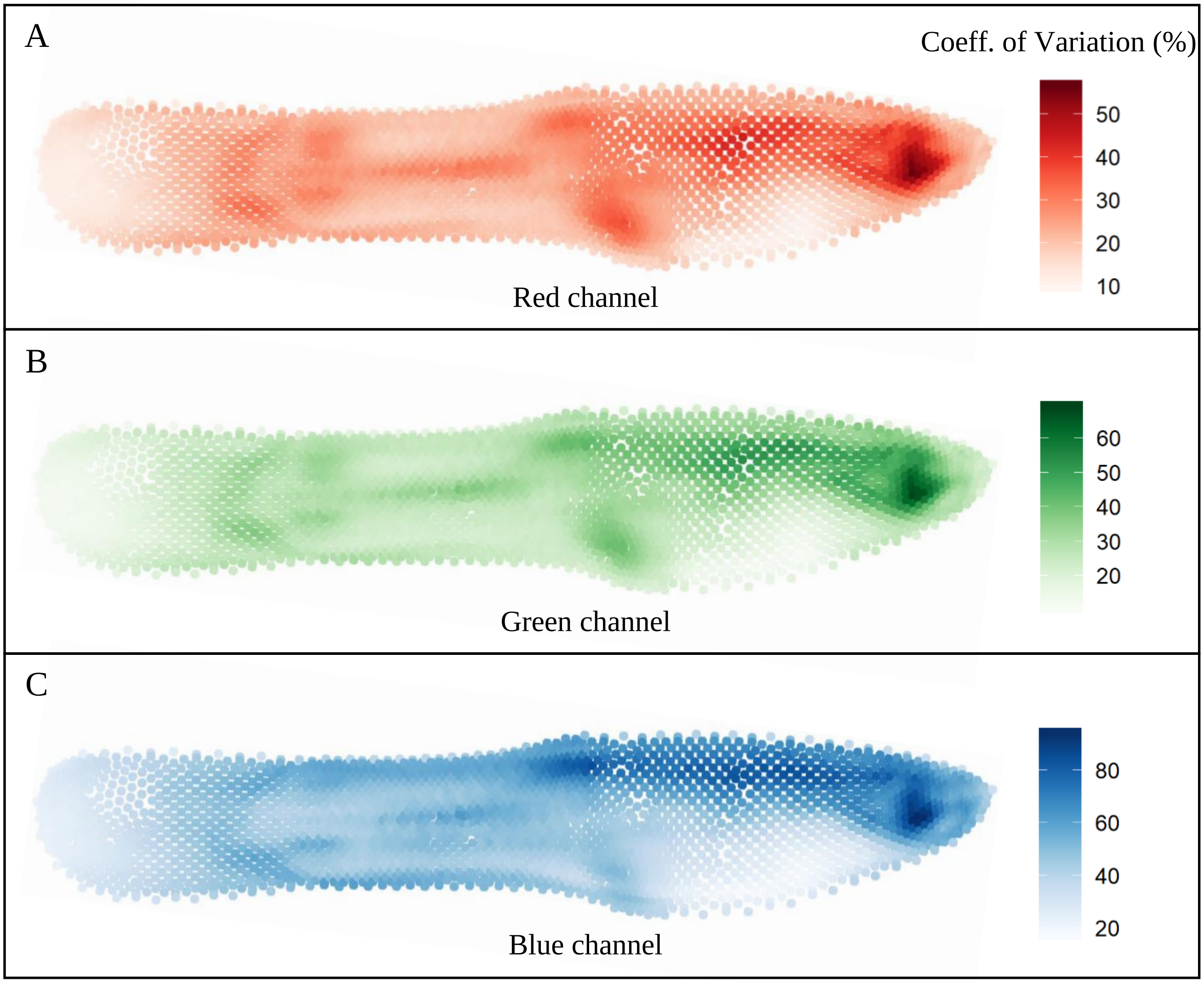


Supplemental Figure S4. A heat map showing the coefficient of variation (%) at each of the 2,462 sampling points for the (A) red, (B) green, and (C) blue color channels. Darker colors indicate greater variation among all guppy images (N=485) in the color channel measure at a given sample point.
